# Molecular epidemiology of Japanese Encephalitis Virus in pig population of Odisha, Assam and Manipur states of India

**DOI:** 10.1101/818070

**Authors:** Ankita Datey, Leichombam Mohindro Singh, Uttam Rajkhowa, Birendra Kumar Prusty, Tanuja Saswat, Prabhudutta Mamidi, Luit Moni Barkalita, Rupam Dutta, K Chandradev Sharma, Dinabandhu Sahoo, Probodh Borah, Sarangthem Indira Devi, Soma Chattopadhyay

## Abstract

Japanese encephalitis virus (JEV) comes under the family *Flaviviridae* and genus flavivirus. It predominantly infects the children under the age of 10 years and the case fatality rate can stretch out as high as 30%. Pigs act as reservoir and amplifying intermediate host for JEV. Recent report suggested longer persistence of JEV in tonsil than in circulation of experimentally infected pigs. The current investigation was conducted to understand the prevalence and molecular epidemiology of JEV infection in pigs in three different geographical sites in India (Odisha, Assam and Manipur). Serum samples were tested by ELISA and RT-PCR for detection of JEV, while only RT-PCR was done in case of tonsils tissues collected from pigs slaughtered in abattoir. Prevalence of JEV was highest in Manipur (25.45% in serum and 10.08% in tonsil) but lower in Assam (3.75% in serum and 0% in tonsils) and Odisha (1.49% in serum and 3.7% in tonsils). The percentage of sero-positivity was found to be 3.75% of IgM and 9.9% of IgG in Assam and Odisha respectively. Genotype III (GIII) of JEV was the dominant genotype and sporadic mutations of S83G, H76P, E78Q, C55S, and S64W along with two consistent mutations V46S and V51I were observed in all the GIII strains. Analysis of the E gene sequence revealed a single mutation, S118N in the GI strain. Older pigs (above 7 months) were found to be infected relatively more (8.6%) than younger pigs (age group 3-7 months). In conclusion, the high JE virus infection rate of pig in the current locations suggests the need for continuous surveillance of this virus in pigs which will ultimately help to adopt an effective control strategy to prevent the spread of JE infection to human.

**Author summary:** Japanese encephalitis is one of the contributing factors in acute encephalitis syndrome cases reported across India as well as Asia. Primarily young naive human population are affected with JEV. The death rate can be as high as 30% and in about 30%-50% surviving population paralysis, brain damage or other serious permanent sequelae may be observed. The viral load gets amplified in pigs and thus plays a crucial role in transmitting the infection in human communities living in close proximity to pig dwelling. The current study was conducted to demonstrate prevalence of JEV in pig population of three geographical regions of India *viz.* the States of Odisha, Assam and Manipur that have reported JE outbreaks in human population. The current study demonstrates that the rate of infection is 3.28% among pigs in Manipur followed by Assam and Odisha. GIII was found to be the most predominant JEV genotype, while only one GI genotype strain was detected from Odisha region. These findings suggested the need of continuous surveillance of this virus in pigs and proper implementation of human and animal vaccination programme to control the infection.

## Introduction

Flaviviruses are important human and animal pathogens with worldwide distribution. Japanese encephalitis virus (JEV), belonging to this family particularly affects the children below 10 years and the mortality rate can be upto 30% [1] and about 30%-50% of survivors develop permanent neurologic disorder or psychiatric sequelae [2]. JEV is a single stranded RNA virus with a genome length of approximately 11kb. The genome is divided into a structural region containing capsid (C), pre-membrane (prM) and envelope (E) genes and a non-structural region consisting of NS1, NS2a, NS2b, NS3, NS4a, NS4b and NS5 genes [3]. The NS3 and NS5 genes encode for viral helicase and polymerase enzymes respectively whereas the functions of NS4a and NS4b are not clearly understood [4]. *Culex tritaeniorhyncus* is the dominant transmitting vector [5] whereas other species of mosquitoes namely *Culex modestus* [6], *Culex pipiens* [7], *Culex bitaeniorhyncus* [7] and *Anopheles sinensis* [8] have also been reported. In nature, JEV maintains its life cycle between vectors and reservoir hosts such as pigs, wading birds and bats [2] from where the virus is transmitted to the dead end hosts such as humans and horses which do not develop high viremia to infect mosquito hosts [9, 10]. Similarly, JEV infection to pregnant sows leads to reproductive failures such as abortions and stillbirths [11]. In South East Asia, JEV transmission is mainly associated with the onset of rainy season but it can occur throughout the year in tropical regions [4].

The JEV strains have been characterized genetically depending upon the limited sequencing of C/prM, E and NS5/3′ UTR regions, while based on their evolutionary divergence JEV genotypes are further categorized into GI (GI-a and GI-b), GII, GIII, GIV and GV [12, 13]. A single serotype exists for all the JEV genotypes [14]. The Nakayama and Beijing-1 prototype strains and all other strains isolated from pigs and mosquitoes before 1994 belong to GIII [15]. However, there are reports indicating recent introduction of GI strain in pigs and mosquitoes [16, 17]. It has also been reported that GI has slowly replaced GIII strain and can become a dominant genotype because of contribution of several viral, environmental and host factors [18]. Moreover, the GI genotype has equal virulence as that of GIII and is responsible for encephalitis in humans in most Asian countries including India [12]. Earlier reports suggested that GIII is the most prevalent genotype in India [19], though there are reports suggesting the presence of GI in Uttar Pradesh [20] and West Bengal [21]. The main cause of viral encephalitis in Asia is JEV. The transmission of JEV is endemic in around 24 countries of South East Asia and Western Pacific regions [22]. More than 3 billion people are at risk of JEV infection with approximately 68,000 clinical cases reported every year [23]. JEV epidemics among human was first reported in Haryana, India in 1990 [24] and a major outbreak was reported from Gorakhpur in Uttar Pradesh, India in 2005 [1]. Subsequent outbreaks also occurred in Malkangiri in 2012 and Manipur in 2016 [4]. Most of the epidemics occurred in the first to third weeks of October in a year [1]. The common clinical manifestations include fever (temperature 38.5°C-40°C), severe headache, convulsions, vomiting and in extreme cases the infections leads to paralysis, coma and death [1]. According to National Vector Borne Diseases Control Programme, 216 deaths were reported out of approximately 1731 cases in 2015.

JE is mainly reported in the rural areas as most of the Indian population relies on agriculture and the agricultural grounds serve as a potent place for mosquito breeding. As the vector remains in close proximity to its reservoir hosts, this significantly increases the risk of JE transmission to human as well as other hosts such as horses [1]. Pigs are the prime suspects as reservoir for disease transmission [25] and the high viral titer in peripheral blood of pigs ensures transmission of infection to susceptible naive young human population through mosquito bite. Reports also suggest, the transmission of virus from infected to uninfected pigs occurring through mosquitoes [26]. Recent reports indicates the vector free transmission of JEV through the oronasal secretion of infected pigs to naive pigs [25].

There are several reports regarding the mosquito studies in the JE endemic regions of India, however studies on reservoir hosts is limited [19]. As investigating the JEV persistence in the pig population is of utmost importance to take the necessary precautionary measures before the onset of fatal situation, this investigation was conducted to study the molecular epidemiology of JEV in the pig population of JE endemic regions of Odisha, Assam and Manipur in India.

## Materials and methods

### Sample collection

Blood and tonsil tissue from slaughtered pigs were collected from the endemic regions of Odisha, Assam and Manipur from August 2017 -March 2019 (Fig 1). The animals used in the study were being processed as part of the normal work in the abattoir. The tissue and blood samples were collected by qualified practicing veterinarians from animals slaughtered in the abattoir as a part of the routine process after obtaining necessary consent for the same.

**Fig 1.**
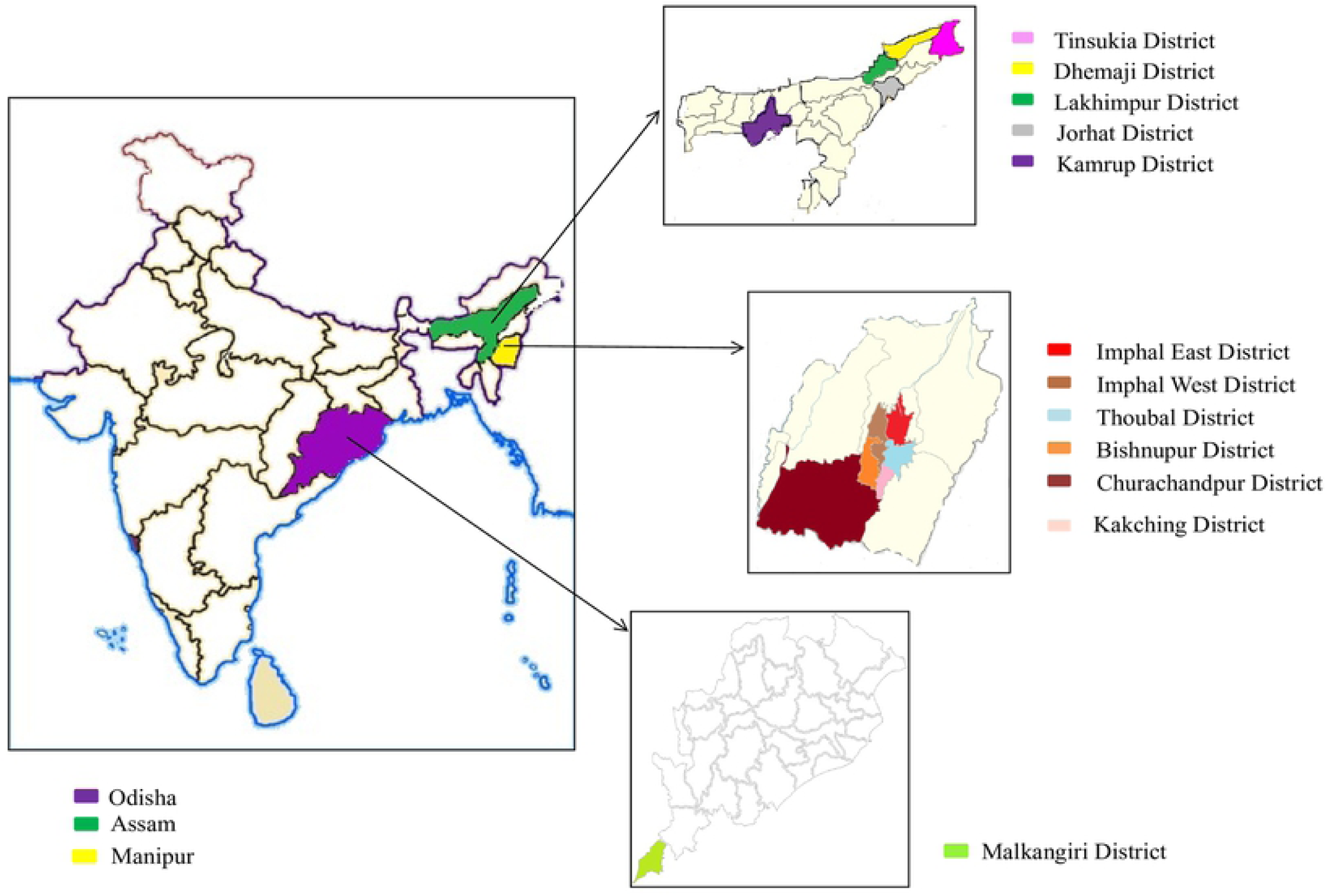
Outline Map of India showing three different States of Odisha, Assam and Manipur. The colored areas represent the districts of different States to focus the exact location from where samples were collected (https://commons.wikimedia.org/wiki/Atlas_of_the_world).

The serum was separated from the blood sample by centrifugation at 5000 rpm for 5 mins at 4°C and aliquoted in properly labeled vials. Tonsil sample of around 1gram was collected from slaughtered pig and stored in RNA later (Ambion, Thermo Fischer Scientific, USA) [16]. Both the serum and the tonsil tissues were stored at −80°C until further processing.

### Viral RNA extraction from serum and RT-PCR

The viral RNA was extracted from serum samples as referred by Desingu *et al*, 2016 [19]. For this, 140μl of swine serum was used and extraction was done, using viral RNA isolation kit (QIAmp Viral RNA Mini Kit Qiagen, Germany) according to the manufacturer’s protocol. The viral RNA was quantified and cDNA was prepared with random hexamers using 1μg of viral RNA by Superscript III first strand cDNA synthesis kit (Invitrogen, USA). This was followed by a PCR reaction using Taq DNA polymerase (Thermoscientific, USA) with initial denaturation at 95°C for 5 mins followed by 35 cycles of 95°C for 45 s, 45°C for 45 s and 72°C for 30 s with the final extension of 10 mins, amplifying 400bp of envelope (E) gene using specific primers [27]. The amplified products were electrophoresed on 2% agarose gel and viewed under UV transilluminator.

### Total RNA extraction from tonsil tissue and RT-PCR

The total RNA was extracted from tonsil tissue as per the method described by Mendez *et al*, 2011 [28]. The tonsil tissue was homogenized in 500μl of TRIzol (Invitrogen, USA) by sterile homogenizer and total RNA was extracted using manual trizol extraction method and quantified. Approximately 1μg of total RNA was used to prepare cDNA with random hexamers using Superscript III first strand cDNA synthesis kit (Invitrogen, USA). The 400bp partial region of E gene was amplified as mentioned earlier.

### Serology

The serum samples were also subjected to indirect ELISA to detect the presence of IgG antibodies against JEV using Porcine JE IgG ELISA Kit (Glory Science Co., Ltd, USA) in accordance with the manufacturer’s instructions.

The IgM antibody detection against JEV in pig serum was also carried out by using Porcine Japanese Encephalitis IgM antibody (JE-IgM AB) ELISA kit (Genexbio Health Science Pvt. Ltd.) as per manufacturer’s protocol.

### Sequencing and phylogenetic analysis using E gene sequence

The positive serum and tissue samples were subjected to bi-directional sequencing with overlapping primers using an automated DNA sequencer (Applied biosystem®3500 series genetic analyser), based on the Sanger’s dye terminating method. The sequence obtained was analyzed for its correct identity using the BLAST tool of NCBI and confirmed.

The obtained sequences were aligned by the ClustalW tool [29] and the phylogenetic analysis was performed using the MEGA6.06 software [30]. The phylogenetic tree was constructed using the Neighbor-Joining method [31] and the evolutionary distances were compared using the maximum composite likelihood method [32]. The statistical significance of the phylogenetic tree was tested with a bootstrap value of 1000 pseudo-replicate datasets [33].

### Mutational analysis

The nucleotide sequences of the JEV E gene of the isolated strains were translated to protein sequences using the ExPASy translate tool (https://web.expasy.org/translate/). The sequences were then aligned with their respective prototype strain using the Clustal Omega tool (https://www.ebi.ac.uk/Tools/msa/clustalo/). The variations in the protein sequences were highlighted.

### Accession numbers

The E gene sequences (n=36) were submitted to GenBank database with accession numbers MK421340, MK940903, MK940904, MK940905, MN115386, MN115387, MK491507, MH376692, MK682387, MK692888, MK692889, MK692890, MK692891, MK692892, MK692893, MK692894, MK518053, MK952776, MK952775, MK962309, MK962310, MK975821, MK975822, MK975823, MK975824, MN029019, MN010526, MN010527, MN010528, MN010529, MN029013, MN029014, MN029015, MN029016, MN029017, MN029018.

## Results

In the present study, the total number of serum samples screened from Odisha, Assam and Manipur were 402, 400 and 55 respectively; whereas the corresponding numbers for tonsil samples were 27, 55 and 248. The number of JEV positive serum samples detected by means of RT-PCR was 6 out of 402 (1.49%), 15 out of 400 (3.75%) and 14 out of 55 (25.45%) and for tissue samples 1 out of 27 (3.7%), 0 out of 55 (0%) and 25 out of 248 (10.08%) in the respective States (Fig 2A and 2B). On the other hand, 15 out of 400 (3.75%) of IgM and 40 out of 402 (9.9%) of IgG positive cases were reported from Assam and Odisha respectively.

**Fig 2.**
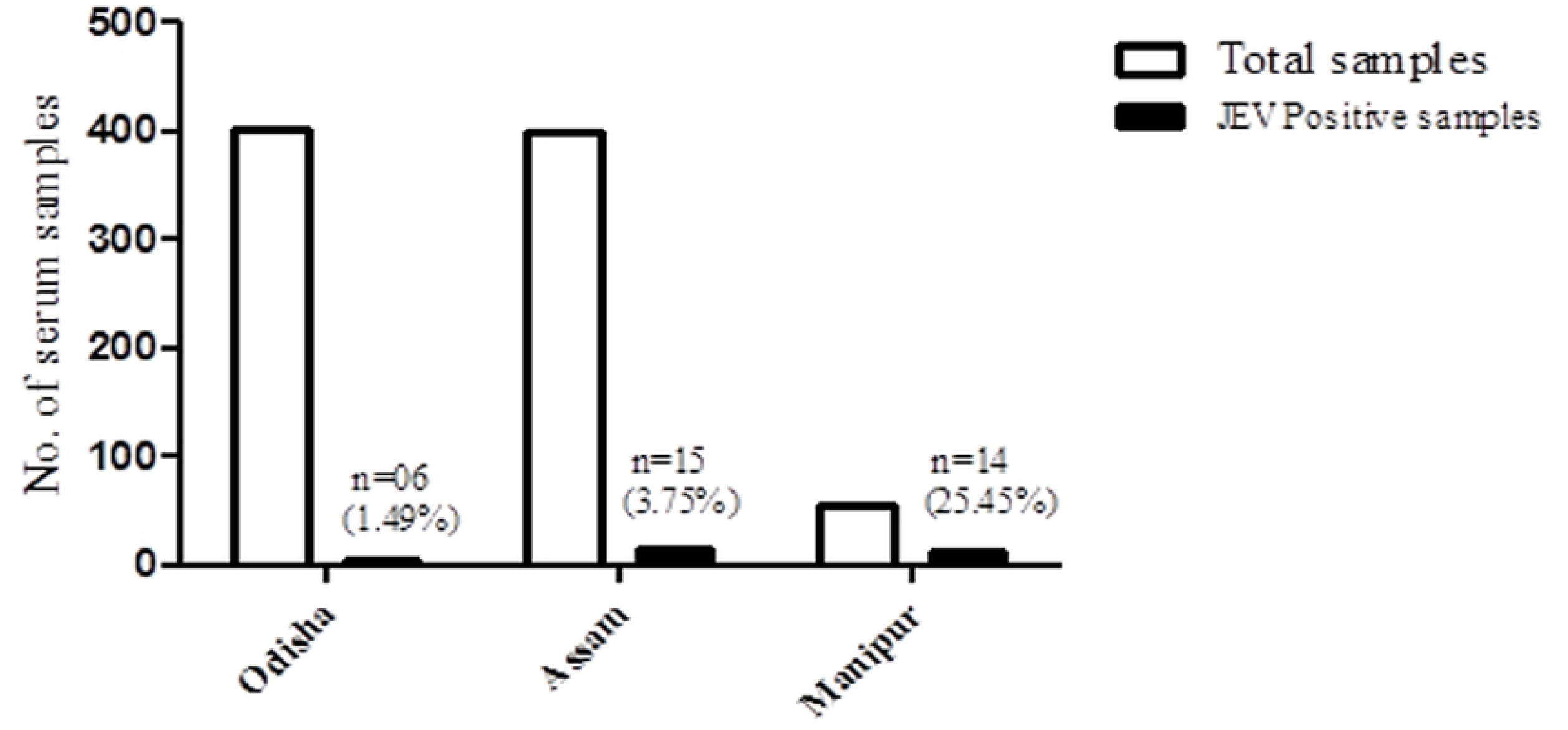
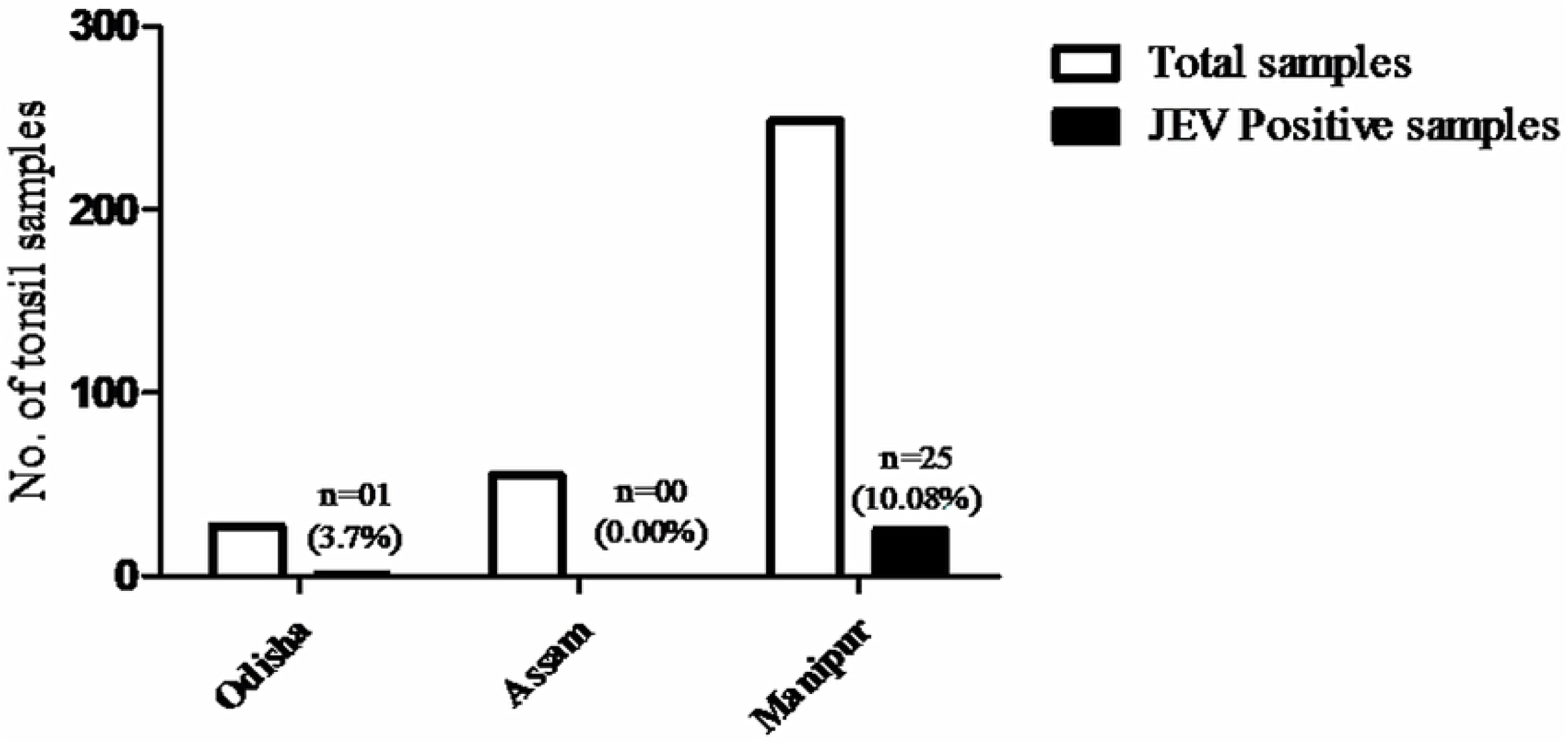
Detection of JEV in samples of pig collected between August 2017-March 2019 by RT-PCR. (A) Number of positive cases from the serum samples collected from pigs. (B) Positive number of cases from the tonsils tissues collected from the slaughtered pigs.

### District-wise distribution of JE positive cases

As per the district-wise distribution for all the three states, the overall prevalence of JEV for serum samples was found to be 1.49% in Malkangiri, 25.45% in Imphal West, whereas 3.33% in Jorhat, 5.71% in Lakhimpur, 4.00% in Dhemaji and 1.67% in Kamrup (S1A Table). Besides this, percentage of positivity for tonsils tissues by RT-PCR were found to be 3.7% in Malkangiri, 8.23% in Imphal West, 35.89% in Imphal East, 3.4% in Kakching and 100% in Bishnupur (S1B Table). However, district wise distribution was not available for Assam tonsil samples. Further, there was not a single positive case for tonsil sample by RT-PCR in Assam (S1B Table).

### Age, month and sex-wise distribution of positive cases of JE in pigs detected by RT-PCR

The number of JEV positive cases detected in pig serum in the above mentioned states was found to be 5.21%, 1.67% and 77.7% respectively for the age group 3-7 months, whereas above 7 months, it was found to be 0%, 6.88% and 15.2%, respectively in Odisha, Assam and Manipur (Fig 3A and 3B). However, for tonsil samples from Manipur 11.12% (3 out of 27) was found to be positive under age group 3-7 months, whereas 9.95% (22 out of 221) was found positive in the age group above 7 months. Details regarding age-wise distribution of positive cases in respect of tonsil samples were not available for Odisha and Assam. In the current study, in Odisha more number of positive cases was found during the pre-monsoon season whereas in Assam, number of positive cases was more during monsoon and post-monsoon periods. The sex-wise distribution of JEV serum positive cases in the three states were 1.85%, 2.56 % and 40.0% in males, and 7.93%, 4.88%, 8.0% in females (Fig 4A and 4B). Similar details in respect of tonsil samples were not known for all the three states.

**Fig 3.**
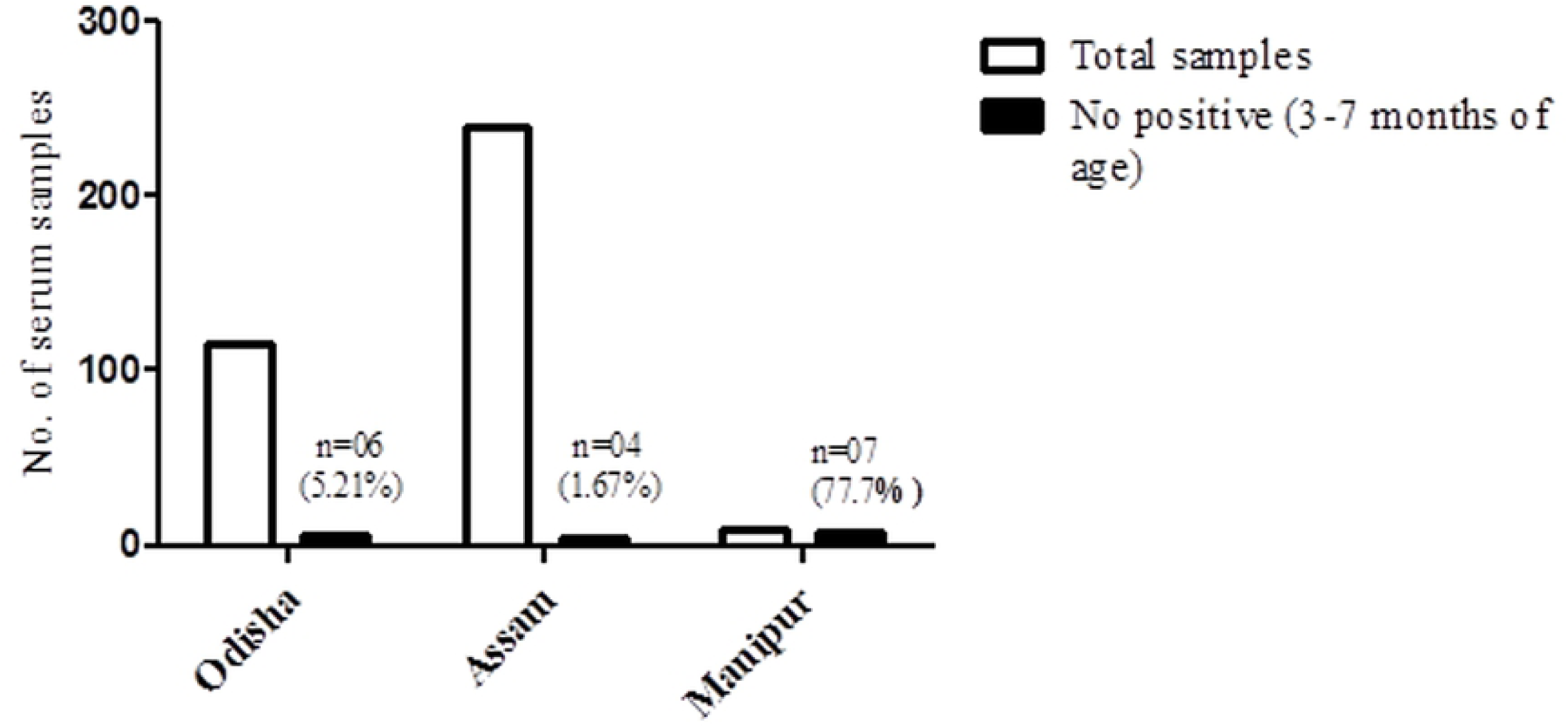
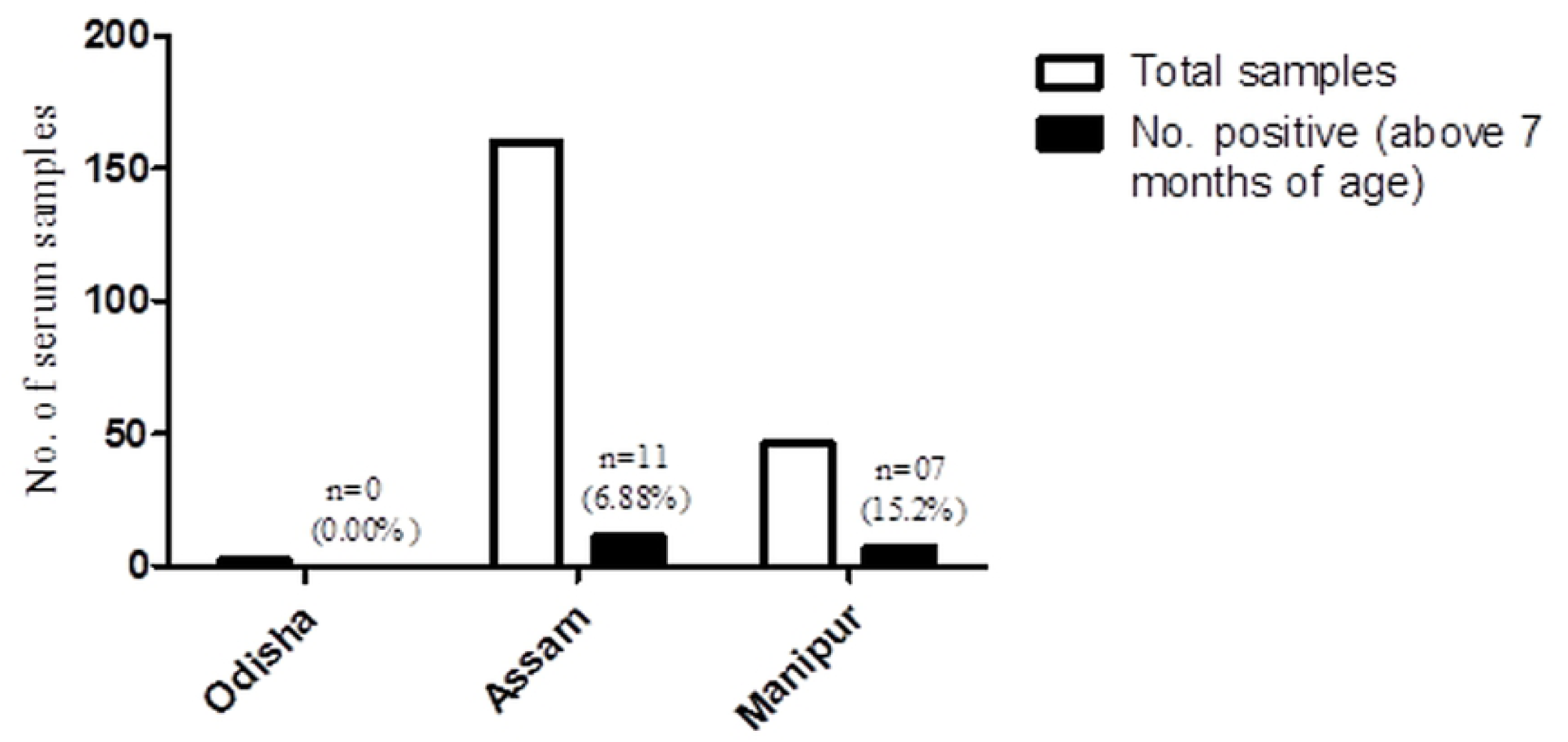
Age-wise distribution of RT-PCR positive JEV cases detected from pig serum. (A) Positive number of cases under age group 3-7 months. (B) Number of positive cases above the age of 7 months

**Fig 4.**
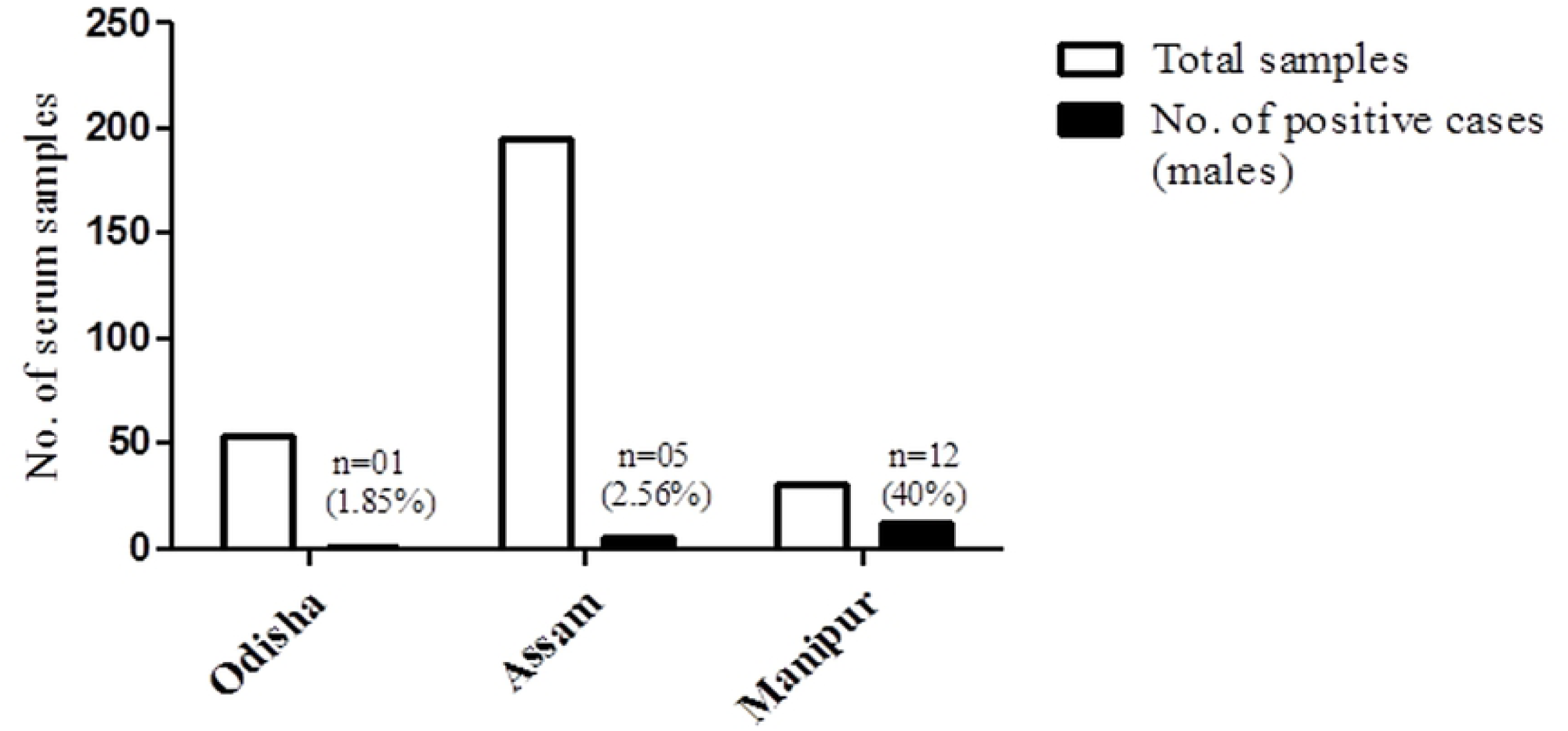
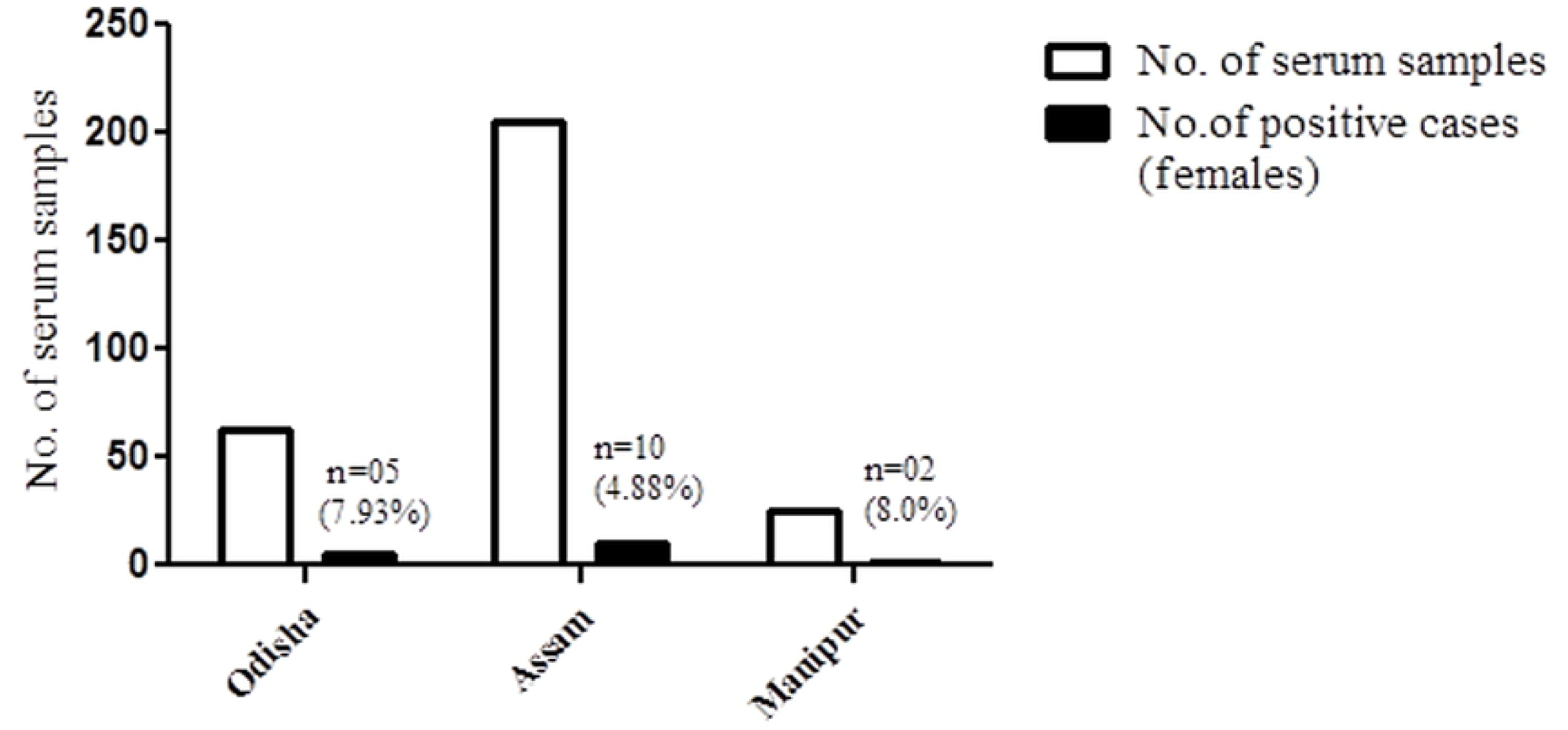
Sex-wise distribution of RT-PCR positive JEV in pig serum. (A)Number of RT-PCR positive male cases. (B) Number of RT-PCR positive females cases.

### Sequencing and phylogenetic analysis

In the present study, a total of 61 samples including serum and tonsil were found to be positive by RT-PCR. The PCR products which were found positive were confirmed by sequencing. Out of 61 positive samples 36 sequences were submitted to the GenBank. All the strains bearing GenBank accession numbers are listed in Table S2.

The genotype I and III strains were found to be circulating among the pigs of the three states under study, of which, genotype III was found to be the predominant one. The genotype III strains (n=60) were found in all the three States whereas a single case of genotype I (n=1) was reported from Malkangiri district of Odisha.

The phylogenetic tree was constructed with reference JEV strains of genotype I (n=19), genotype II (n=2), genotype III (n=6), genotype IV (n=2) and genotype V (n=2) including the strains isolated in the present study. All of the isolated strains in the present study were clustered into genotype I and genotype III. The isolated strain with accession number MK421340 clustered with the other GI strains reported from Vietnam, India, Korea, China, Taiwan and Japan (Fig 5). However, the rest of the strains clustered in genotype III including the prototype Nakayama strain.

**Fig 5.**
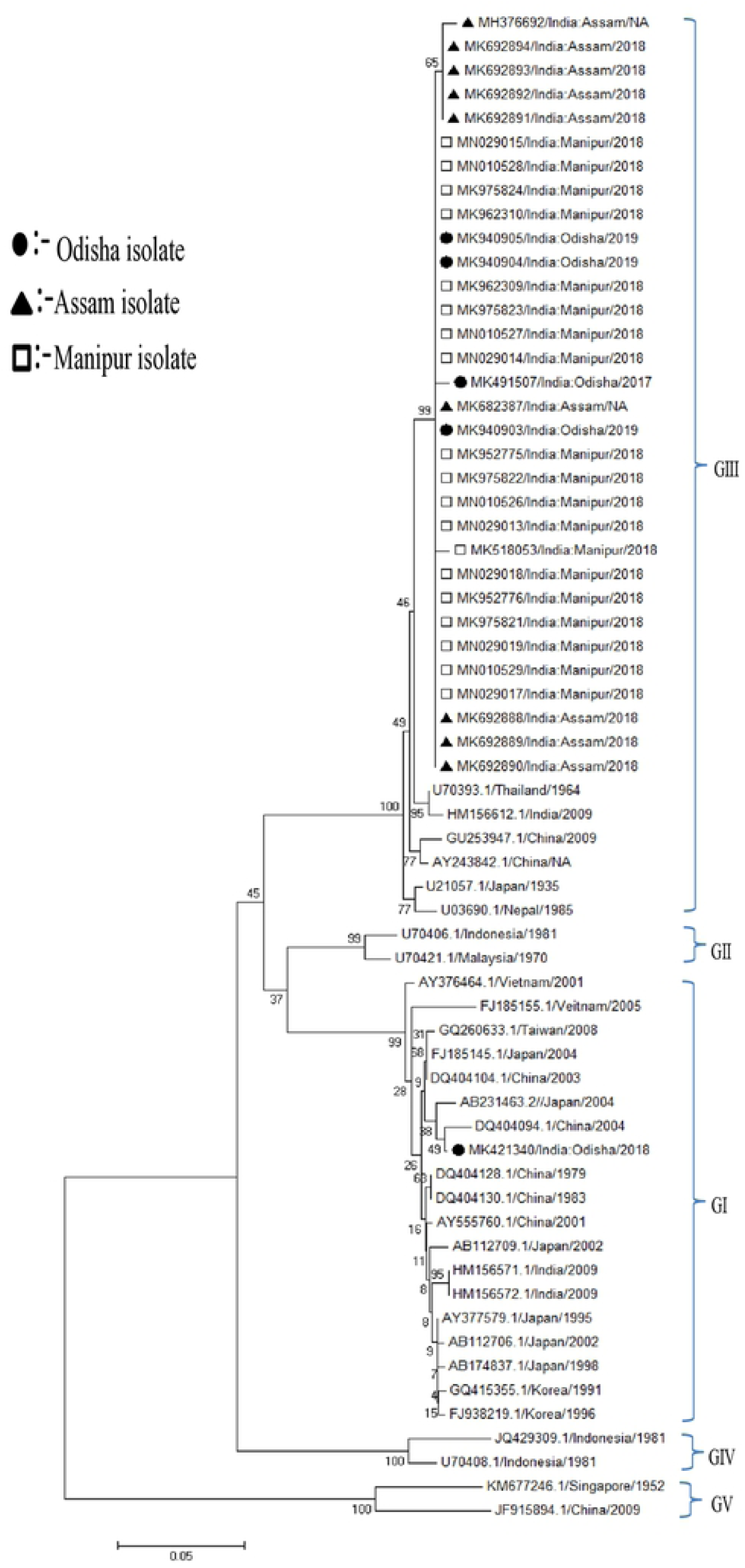
The phylogenetic analysis of the E gene of JEV strains from pigs. The phylogenetic tree was constructed using neighbor-joining method with 1000 bootstrap value by the MEGA6.06 software. The reference strains of GI, GII, GIII, GIV and GV were obtained from the GenBank database. Viruses are depicted by accession number/country/year of isolation. The close circle, close triangle and open square represents the Odisha, Assam and Manipur isolates respectively. The bootstrap values are indicated at major branch points and scale bar indicates nucleotide substitution per sites.

### Presence of mutations in the isolated strains

The amino acid sequence of the translated protein coded by the E gene of the detected GI strain was aligned with that of the respective prototype strain as well as to the strain isolated from human in 2005 from Gorakhpur, India using the Clustal Omega tool. It was noticed that the JEV GI strain isolated from Odisha was approximately 99% identical with a single mutation (S118N) observed within the amino acid region of 1-156 of the E gene (S3A Table and S1A Fig). Similarly, after the alignment of GIII E amino acid sequences with the prototype strain, it was observed that they were approximately 96-98% identical. The sequence analysis showed the presence of two consistent mutations V46S and V51I in all the GIII strains of the present study. Further, S83G mutation was observed in five Assam strains, two mutations S64W and H76P were observed in strain with accession number MH376692, whereas a single mutation E78Q was observed in Odisha strain (MK491507) and C55S mutation was found in Manipur strain (MK518053) (S3B Table and S1B Fig).

## Discussion

Pigs act as an amplifier host of JEV, thus can be a key suspect of spreading infection to human living in close proximity [4]. The detection of JEV in pigs was reported earlier from Cambodia, Singapore, Vietnam, Australia, Japan, China, Thailand, Laos, Indonesia, India etc (Fig 6) [22, 34-42], however the studies on pigs for the presence of JE virus and its genotypes in circulation among pigs in India are limited [19]. Hence, the current investigation was mainly focused on the JE virus detection in pig serum and tonsil in three different JE endemic states of India namely Odisha, Assam and Manipur.

**Fig 6.**
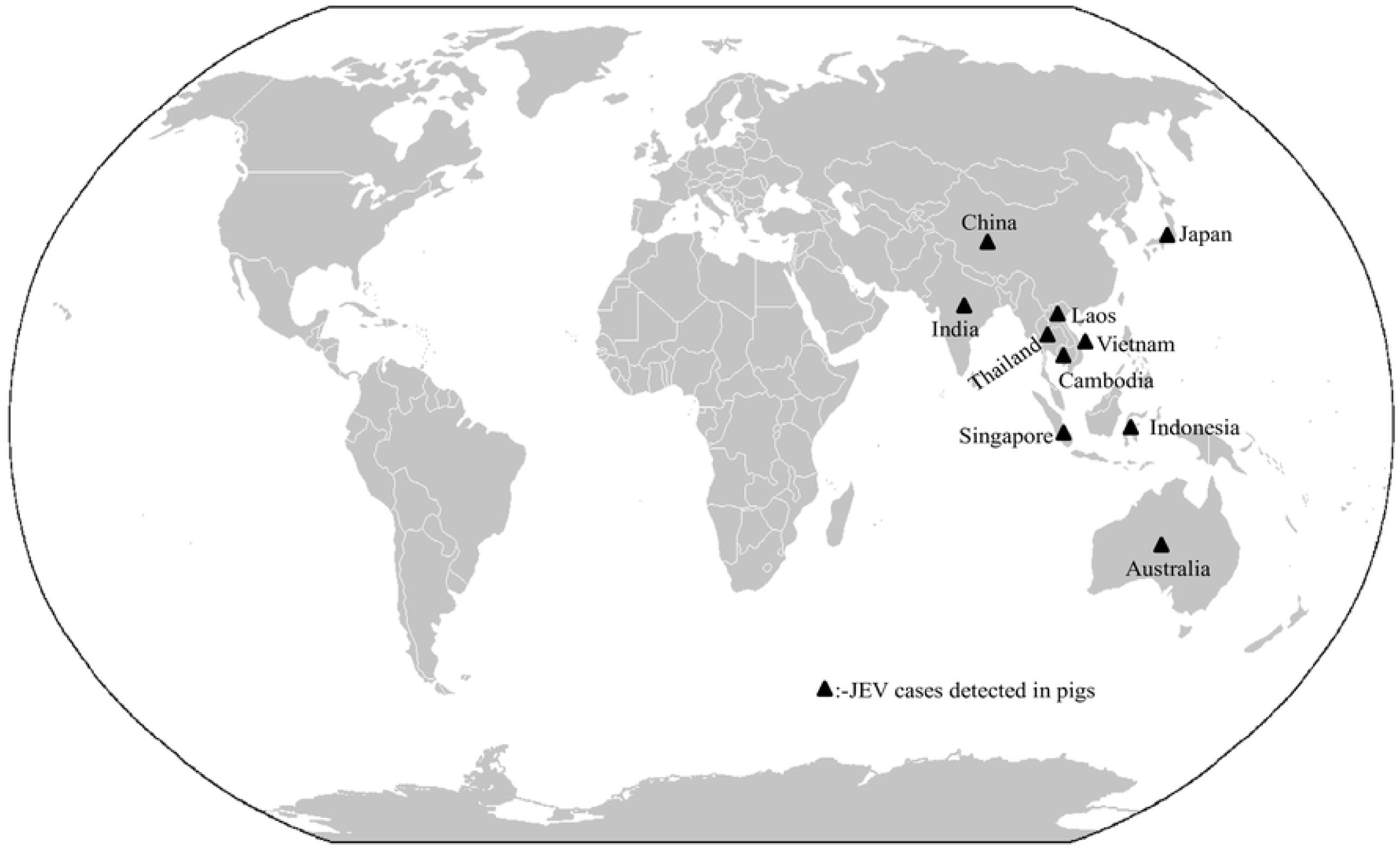
The World map showing reported cases of JEV detection in pig population (https://commons.wikimedia.org/wiki/Atlas_of_the_world).

In the current investigation, a large number of JE positive cases were detected among pigs in Manipur State of India, as compared to the other two states. Moreover, it was noticed that the genotype III strain was predominant in all the three states with a single case of genotype I in Odisha.

In the present study, high sero-positivity was observed in pigs from the districts of Odisha, Assam and Manipur. There are reports that *Culex tritaeniorhynchus* acts as a transmitting vector and majorly found around the pig farming areas throughout the year. There are reports which suggest that the pigs are the potential feeding hosts for *Culex tritaeniorhynchus* [43, 44]. The presence of virus in tissue sample suggests that the virus can also invade tonsil tissue along with CNS as was reported earlier [25]. According to previous reports, in experimental JEV infected pigs, the viral load was noticed to be highest in tonsils as compared to other secondary lymphoid organs as well as CNS. In tonsils, even after 11 days of post infection no significant reduction was observed in viral RNA. Further, live virus was also detected in tonsil tissues [25].

The higher percentage of positive cases detected in Manipur (Imphal West district) and Assam (Lakhimpur district) might be due to larger pig population of the north eastern regions of India [45]. The male pigs were found to be infected more with JE virus compared to females. Moreover, the rate of infection was higher in the pigs above 7 months of age compared to 3-7 months. Pigs over 11 months of age were also found positive for JEV in Cambodia [46]. It might be due to pig to pig transmission through oronasal route in pig rearing farms [25]. The JE positivity was observed throughout the year in the current study with slight increase during the monsoon and post-monsoon periods. Earlier reports also suggested that the JE detection goes up during the monsoon and the virus can be detected in pig throughout the year in tropical regions [4].

Here, the genotype III strain was found to be circulating predominantly among pig population of the three states. This was reported earlier that genotype III is the most prevalent strain in pig population of India as well as other parts of Asia such as China [19, 47]. Interestingly, a single strain of genotype I was detected in pig population of Odisha. There are reports which indicate the recent introduction of genotype I in human in India [20], however GI JEV strain has never been detected previously from pig. The presence of genotype I was also reported from other parts of Asia, suggesting it as an emerging strain [15]. The highest nucleotide identity for the GenBank submitted sequence of genotype I (GenBank Accession number MK421340) was found to be 99.74% with Yunan isolate YNTC07290. Whereas, for genotype III the submitted sequences of GenBank showed highest nucleotide identity of approximately 99% with Indian isolate JEV/SW/IVRI/395A/2014. As reported earlier also, genetic diversity is low for the E gene at nucleotide and amino acid level [23]. The mutational analysis showed a single mutation in GI strain, whereas sporadic mutations along with two consistent mutations were observed for all the GIII strains.

In the present investigation, paired samples of serum and tonsil could not be collected due to operational issues in the abattoir, which could have offered better insights into the merit of testing tonsil tissue (if any) rather than serum samples for detection/quantification of JEV by RT-PCR. Further, a well-designed survey for JEV infection/disease prevalence in human and mosquito population during the study period would have allowed possibilities to correlate JEV infection in humans and pigs in the same geographical region.

The current study limited to three geographical sites has indicated that JEV prevalence in pigs can be very high in areas where human JEV outbreak has been reported. We suggest that JEV outbreaks in human population might be controlled by vaccinating pigs in areas of high prevalence of the viral load in the animal reservoir – a strategy that has not been widely adopted.

## Acknowledgement

We are grateful to Dr Anirban Basu (NBRC, Haryana, India) for kindly providing the JEV strain, GP78. We are thankful to Dr Balachandran Ravindran for conceptualization, reviewing and editing the manuscript. We would like to thank Ujjwal Mahata and BirSingh Mahata for collecting the pig samples from Malkangiri region of Odisha. We are also grateful to Konthoujam Abung Meitei for providing samples from Imphal West district of Manipur.

## Supporting information captions

S1A Table. District-wise distribution of the positive JEV cases by RT-PCR from the pig serum samples collected during the study period. (DOC)

S1B Table. District-wise distribution of the positive JEV cases by RT-PCR from the pig tonsil samples collected during the study period. (DOC)

S2 Table. Accession numbers for the positive serum and tonsil samples from three different States of India. (DOC)

S3A Table. Amino acid substitutions in the envelope (E) gene of the studied JEV genotype I (GI) strain in comparison to the prototype strain. (DOC)

S3B Table. Amino acid substitutions in the envelope (E) gene of the studied JEV genotype III (GIII) strains in comparison to the prototype Nakayama strain. (DOC)

S1A Fig. Alignments of JEV E amino acid sequences of GI strain detected in the study in comparison to strains isolated from mosquito and human from China and India respectively. Conserved amino acids are kept as star marked. The substitution mutations are represented in colour with highlight (DOC).

S1B Fig. Alignments of JEV E amino acid sequences of GIII strain detected in the study in comparison to prototype Nakayama strain. Conserved amino acids are kept as star marked. The substitution mutations are represented in colour with highlight. (DOC)

